# Pharmacological inhibition of GPR4 remediates intestinal inflammation in a mouse colitis model

**DOI:** 10.1101/533174

**Authors:** Edward J. Sanderlin, Mona Marie, Juraj Velcicky, Pius Loetscher, Li V. Yang

## Abstract

Inflammatory bowel disease (IBD) is characterized by chronic, recurring inflammation of the digestive tract. Current therapeutic approaches are limited and include biologics and steroids such as anti-TNFα monoclonal antibodies and corticosteroids, respectively. Significant adverse drug effects can occur for chronic usage and include increased risk of infection in some patients. GPR4, a pH-sensing G protein-coupled receptor, has recently emerged as a potential therapeutic target for intestinal inflammation. We have assessed the effects of a GPR4 antagonist, 2-(4-((2-Ethyl-5,7-dimethylpyrazolo[1,5-a]pyrimidin-3-yl)methyl)phenyl)-5-(piperidin-4-yl)-1,3,4-oxadiazole (GPR4 antagonist 13, also known as NE 52-QQ57) in the dextran sulfate sodium (DSS)-induced acute colitis mouse model. The GPR4 antagonist 13 inhibited intestinal inflammation. The clinical parameters such as body weight loss and fecal score were reduced in the GPR4 antagonist 13 treatment group compared to vehicle control. Macroscopic disease indicators such as colon shortening, splenic expansion, and mesenteric lymph node enlargement were all reduced in severity in the GPR4 antagonist 13 treated mice. Histopathological features of active colitis were alleviated in GPR4 antagonist 13 treatment groups compared to vehicle control. Finally, inflammatory gene expression in the colon tissues and vascular adhesion molecule expression in the intestinal endothelia were attenuated by GPR4 antagonist 13. Our results indicate that GPR4 antagonist 13 provides a protective effect in the DSS-induced acute colitis mouse model, and inhibition of GPR4 can be explored as a novel anti-inflammatory approach.

## 1. Introduction

Inflammatory bowel disease (IBD) is complex and characterized by dysregulated mucosal immune responses. Uncontrolled chronic gastrointestinal inflammation can result in severe complications which include fistulas and stenosis (Hendrickson et al., 2002; Kaser et al., 2010). Furthermore, IBD can have extrainstestinal manifestations such as arthritis and uveitis (Hendrickson et al., 2002). Additionally, IBD can also increase the risk of the development of colon cancer (Mattar et al., 2011). The current therapeutic landscape for IBD is limited and predominately focused on symptomatic management (Neurath, 2017).

IBD can take the form of ulcerative colitis (UC) and Crohn’s disease (CrD). One feature noted in IBD is the display of reduced intra-intestinal luminal pH in IBD patients compared to healthy controls (Fallingborg et al., 1993; Nugent et al., 2001). It is also well documented local tissue pH can be reduced during active inflammation, predominately owing to heightened metabolic byproducts from excessive leukocyte infiltration and hypoxia (Justus et al., 2015; Lardner, 2001; Okajima, 2013). These studies collectively suggest the intestinal mucosa and lumen can be acidic in IBD patients. It remains poorly investigated, however, how cellular constituents involved in the mucosal immune response sense altered pH in the intestine and subsequently alter function.

G-protein-coupled receptors (GPCRs) are implicated in normal intestinal function as well as in pathological contribution during active IBD. One such family of GPCRs are the proton-sensing GPCRs, which consist of GPR4, GPR65 (TDAG8), and GPR68 (OGR1) (Justus et al., 2013; Justus et al., 2017; Ludwig et al., 2003; Sanderlin et al., 2015). These receptors have been implicated in the modulation of intestinal inflammation (de Valliere et al., 2015; Lassen et al., 2016; Sanderlin et al., 2017; Wang et al., 2018). Studies have shown GPR4 is responsible for acidosis-induced endothelial cell inflammation and can functionally increase leukocyte adhesion with endothelial cells (Chen et al., 2011; Dong et al., 2017; Dong et al., 2013; Tobo et al., 2015). GPR4 genetic deletion in mice reduces intestinal inflammation and mucosal leukocyte infiltration in both inducible and spontaneous colitis mouse models (Sanderlin et al., 2017; Wang et al., 2018). However, the effects of pharmacological modulators of proton-sensing GPCRs in intestinal inflammation have not been investigated.

In this study, we have assessed a GPR4 antagonist, 2-(4-((2-Ethyl-5,7-dimethylpyrazolo[1,5-a]pyrimidin-3-yl)methyl)phenyl)-5-(piperidin-4-yl)-1,3,4-oxadiazole (GPR4 antagonist 13, also known as NE 52-QQ57), within the dextran sulfate sodium (DSS)-induced acute colitis mouse model as a potential therapeutic for the remediation of intestinal inflammation. GPR4 antagonist 13 was recently developed by Novartis and shown effective following oral administration (Velcicky et al., 2017). GPR4 antagonist 13 could reduce arthritis inflammation, angiogenesis, and nociception in animal models (Velcicky et al., 2017). Furthermore, oral pharmacokinetics, GPR4 target selectivity, dosage and potency was thoroughly evaluated (Velcicky et al., 2017). Another study evaluated GPR4 antagonist 13 in mouse ventilatory responses and observed no obvious toxicities (Hosford et al., 2018). Our results demonstrated that GPR4 antagonist 13 administration reduced disease severity, histopathological features, inflammatory gene expression, and endothelial-specific adhesion molecule expression within the inflamed intestinal tissue in the DSS-induced colitis mouse model.

## 2. Materials and methods

### 2.1. Dextran sulfate sodium (DSS)-induced colitis mouse model and GPR4 antagonist 13 delivery

Male and female mice of ages 9-10 weeks in the C57BL/6 background were used for experiments. As previously described, mice were maintained specific pathogen free and housed in an Association for Assessment and Accreditation of Laboratory Animal Care (AAALAC)-accredited facility (Sanderlin et al., 2017). The animal use protocol was approved by the Institutional Animal Care and Use Committee (IACUC) of East Carolina University. Acute experimental colitis was induced by the addition of 3% (w/v) Dextran Sulfate Sodium Salt (DSS) *ad libitum* into the normal drinking water of mice (36,000–50,000 M.Wt, Lot# Q5229, MP Biomedical, Solon, OH). Mice were provided normal pelleted diet *ad libitum* during experimentation (ProLab 2000, Purina Mills, St. Louis, MO). Mice were treated for seven days with 3% DSS with replacement of the 3% DSS solution every two days. As previously described, during each day of the experimental course clinical parameters for disease severity were assessed (Sanderlin et al., 2017). Mice were weighed for body weight loss evaluation and feces was assessed for fecal blood and diarrhea using the Hemoccult Single Slides screening test (Beckman Coulter, Brea, CA). For GPR4 antagonist 13 administration, the small molecule was suspended in 0.5% methylcellulose/ 0.5% Tween 80/ 99% water and a dosage of 30mg/kg (b.i.d.) was given to mice as previously described (Velcicky et al., 2017). On day one, mice were orally gavaged with either vehicle or 30mg/kg GPR4 antagonist 13 in the morning followed by addition of 3% DSS into the drinking water and another dose of GPR4 antagonist 13 or vehicle in the afternoon. On days two through six, mice were orally gavaged with vehicle or GPR4 antagonist 13 twice a day (b.i.d.). On day seven mice were euthanized for tissue collection and macroscopic disease indicator measurement (Figure 1A). Mice were dissected as previously described and mesenteric lymph node expansion, colon shortening, and splenic expansion were measured (Sanderlin et al., 2017).

**Figure 1:**
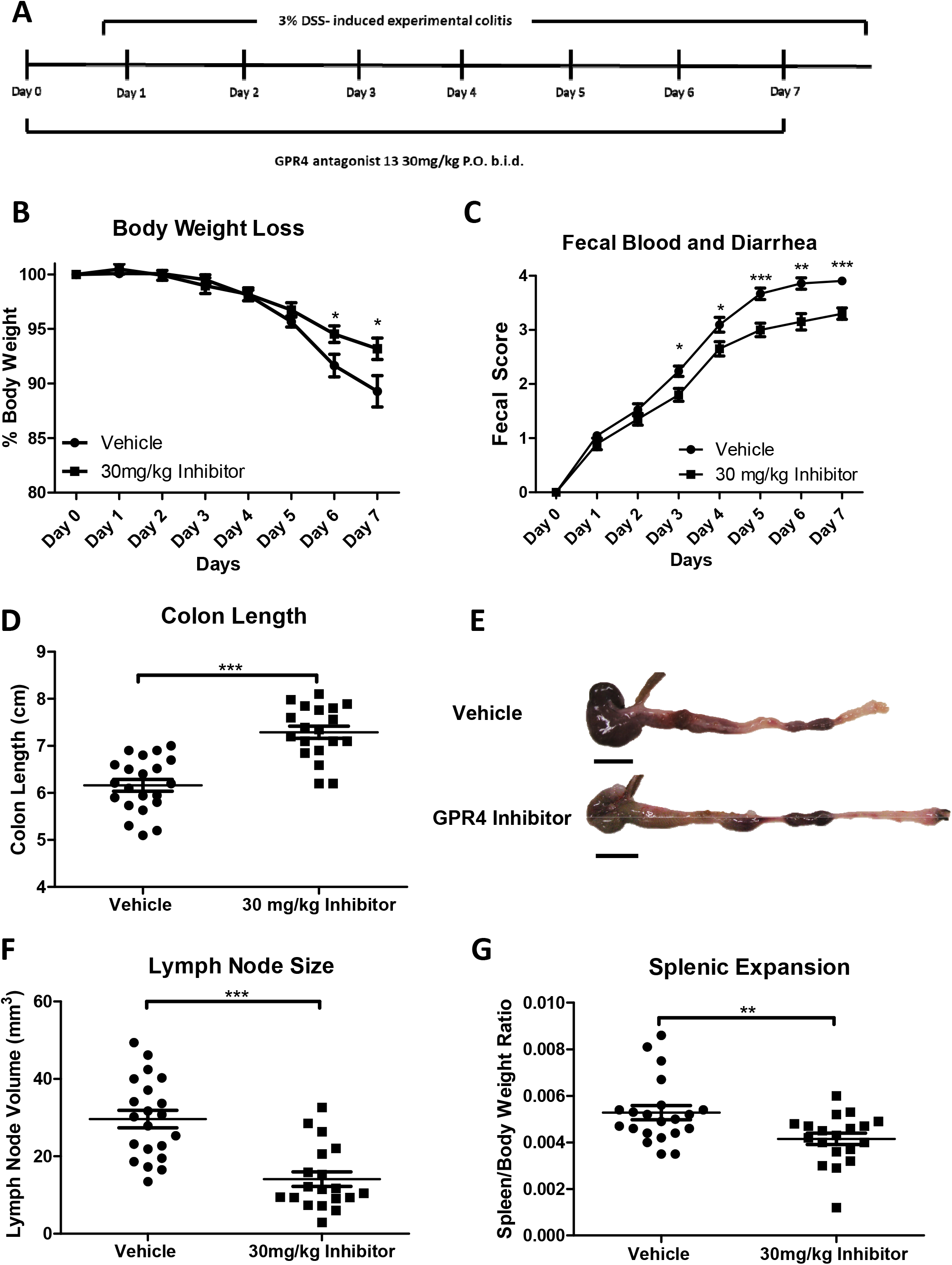
GPR4 antagonist 13 reduces clinical severity and macroscopic disease indicators of intestinal inflammation in mice. Mice were provided GPR4 antagonist 13 P.O. twice a day (b.i.d.) during experimental time course (A). Mouse body weight loss (B) and fecal blood and diarrhea scores (C) were daily measured. Mouse colon length (D-E), mesenteric lymph node expansion (F), and splenic enlargement (G) were also assessed upon tissue collection. Vehicle: N=21 (10 male/11 female) and GPR4 antagonist 13: N=19 (10male/9 female). Data are presented as mean ± SEM and was analyzed for statistical significance using the t-test between vehicle and GPR4 antagonist 13 groups. (*P < 0.05, **P < 0.01, *** P < 0.001). Scale bar = 1cm.

### 2.2. Sample collection for histology and molecular analysis

Following the completion of the seven-day DSS treatment, mice were euthanized and tissue was collected for histology and molecular analysis. The spleen was resected and weighed for splenic expansion. The mesenteric lymph nodes were identified using a dissection microscope, measured for volume calculation, and stored in 10% buffered formalin as previously described (Sanderlin et al., 2017). The gastrointestinal tract was then removed from the mouse and the colon was separated from the cecum at the ileocecal junction. Next, the colon content was removed by PBS flushing and opened along the anti-mesenteric border. Colon tissue approximately 1mm in width was removed from the anus to proximal colon. This tissue was then cut into thirds for distal, middle, and proximal colon segments and promptly snap frozen in liquid nitrogen for molecular analysis. The remaining colon segment was fixed in 10% buffered formalin (VWR) and cut into thirds as distal, middle, and proximal colon segments for histological analysis.

### 2.3 Histopathological analysis

Distal, middle, and proximal colon segments were fixed in 10% buffered formalin, embedded in paraffin and sectioned at 5μm thickness. Slides were then stained with hematoxylin and eosin (H&E) for histopathological analysis. Colon segments were evaluated by operators blind to sample identification according to previously published criteria with minor modifications (Erben et al., 2014). Briefly, each colon segment was evaluated in four recurring locations. Each location was evaluated for leukocyte infiltration, epithelial damage, and mucosal architecture distortions. The leukocyte infiltration score included severity scores of 1= mild, 2= moderate, and 3= severe with regard to both degree and location of cellular infiltrates. The mucosal architecture score included 1= focal epithelial erosions, 2= focal ulcerations, and 3= extended ulcerations. The sum score of these parameters represents the histopathological score of severity.

### 2.4. RNA collection and real time RT-PCR

Total RNA was collected from the distal colon segments using the IBI Scientific DNA/RNA/Protein extraction Kit (MidSci, St. Louis, MO). 500ng of RNA was reversed transcribed using SuperScript IV and diluted 1:10 in ultrapure water (Gibco, Gaithersburg, MD). TaqMan pre-designed primer-probe sets used for the evaluation of specific gene expression (Applied Biosystems, Waltham, MA). TaqMan primer-probe sets and subsequent assay ID include 18S rRNA; (Hs99999901_s1), PTGS2; (Mm00478374_m1), VCAM-1; (Mm01320970_m1), MAdCAM-1; (Mm00522088_m1), E-selectin; (Mm00441278_m1), IL1-β; (Mm00434228_m1), TNF-α; (Mm00443258_m1), IL-10; (Mm01288386_m1), and IL-6; (Mm00446190_m1). Real-time PCR was performed in duplicate with a program of 50°C for 2 min, 95°C for 10 min followed by 40 cycles of 95°C for 15 sec and 60°C for 1 min. Data was acquired using the QuantStudio 3 Real-Time PCR system and analyzed using the 2^−ΔCt^ method.

### 2.5. Immunohistochemistry

Immunohistochemistry (IHC) was performed as previously described (Sanderlin et al., 2017). Briefly, serial 5μm paraffin-embedded sections of distal, middle, and proximal colon segments were deparaffinized and hydrated to water. Antigen retrieval was then performed using Tris-EDTA pH 9.0 with 0.1% Tween 20. Following endogenous peroxidase blocking and AVIDIN/BIOTIN block (Invitrogen, Carlsbad, CA), tissue sections were incubated with primary antibodies against VCAM-1 (Abcam, ab134047, 1:100, Cambridge, MA), MAdCAM-1 (Abcam, ab90680, 1:500, Cambridge, MA) or E-selectin/CD62E (Abcam, ab18981, 1:1000, Cambridge, MA) overnight at 4°C. The rat or rabbit VECTASTAIN Elite ABC HRP kit was then used according to the manufactures protocol (Vector Laboratories, Burlingame, CA). The ImmPACT DAB (Vector Laboratories, Burlingame, CA) substrate was utilized for signal detection. Slides were then dehydrated and mounted. Pictures were taken with the Zeiss AxioImager.M2 with Axiocam 503 digital color camera.

### 2.6. Blood vessel VCAM-1 and E-selectin intensity score and MAdCAM-1 positive vessel enumeration

After IHC was performed for VCAM-1 and E-selectin on colon tissue segments, VCAM-1 and E-selectin intensity was blindly assessed by two independent operators from distal, middle, and proximal segments. Scoring criteria included 1= none/minimal, 2= mild, 3= moderate, and 4= high signal intensity. Each tissue segment was completely evaluated using 10x and 20x objectives and subsequently scored for intensity. For MAdCAM-1 positive vessel enumeration, vessels were assessed from the distal, middle, and proximal colon segments and total MAdCAM-1 positive vessels were counted using 10x and 20x objectives. The total colon length was recorded in centimeters and results were shown as MAdCAM-1 positive vessels/centimeter.

### 2.7. Statistical analysis

GraphPad Prism software was utilized for all statistical analysis. Results were recorded as the mean ± standard error. When analysis was performed between two groups the unpaired *t* test or Mann-Whitney test was utilized. All comparisons *P* < 0.05 are considered statistically significant where * *P* < 0.05, ** *P* < 0.01, and *** *P*< 0.001.

## 3. Results

### 3.1. GPR4 antagonist 13 reduces the severity of colitis in the acute DSS-induced colitis mouse model

Wild-type C57BL/6 mice were initiated on the acute DSS colitis model and given vehicle or 30mg/kg of GPR4 antagonist 13 b.i.d. by oral gavage (Figure 1A). During each day of the experimental course mouse body weight in conjunction with fecal blood and diarrhea scores were evaluated to provide clinical assessment of disease severity between vehicle and GPR4 antagonist 13 groups. GPR4 antagonist 13 treated mice were protected from body weight loss commencing from day five through seven (Figure 1B). Mice given vehicle lost 11-14% body weight by day seven while mice provided GPR4 antagonist 13 lost 5-6% body weight. Fecal blood and diarrhea scores provided further indication GPR4 antagonist 13 protects against intestinal inflammation as mouse fecal scores were reduced in GPR4 antagonist 13 treated mice compared to vehicle (Figure 1C). Fecal blood and diarrhea could be observed in mice with progressive severity from day one through seven. Vehicle mice developed heighted fecal scores earlier than GPR4 antagonist 13 treated mice and maintained more severe scores throughout the seven-day DSS experiment. These results collectively provide evidence GPR4 antagonist 13 can blunt the clinical severity of DSS-induced intestinal inflammation.

### 3.2. Clinical severity and macroscopic disease indicators are reduced in mice treated with GPR4 antagonist 13

Upon completion of the seven DSS-induced experimental colitis, mice were dissected for assessment of macroscopic disease indicators such as colon shortening, mesenteric lymph node expansion, and splenic expansion. We observed that mice treated with GPR4 antagonist 13 had reduced colon shortening when compared to vehicle (colon length ~7.3cm versus ~6.1cm, respectively) (Figure 1D-E). Mesenteric lymph nodes (MLNs) were also collected, and the volume was measured in vehicle and GPR4 antagonist 13 treated mice. Vehicle MLN volume was expanded by more than 2-fold when compared to GPR4 antagonist 13 mice (Figure 1F). These results indicate GPR4 antagonist 13 reduces colonic inflammation and associated expansion of MLNs when compared to vehicle. Finally, the spleen to body weight ratio was assessed in vehicle and GPR4 antagonist 13 DSS-treated mice. We observed reduced splenic expansion in mice treated with GPR4 antagonist 13 when compared to vehicle suggesting reduced disease severity due to GPR4 inhibition (Figure 1G).

### 3.3. GPR4 inhibition reduces histopathological features in mouse colon

To assess the effects of GPR4 inhibition at the histological level, distinct pathological cellular features of colitis were assessed in the distal, middle, and proximal colon segments. Some such features assessed in the colon were leukocyte infiltration, epithelium erosion, crypt distortion, and mucosal ulceration. In both vehicle and GPR4 antagonist 13 DSS-treated mice, the highest degree of histopathology can be observed in the distal colon segments followed by the middle and proximal, respectively. The observation that intestinal inflammation is most severe in the distal colon within the DSS model are consistent with literature (Chassaing et al., 2014; Kim et al., 2012; Laroui et al., 2012; Perse and Cerar, 2012). When comparing vehicle and GPR4 antagonist 13 mouse groups, the degree of histopathology was significantly reduced by GPR4 antagonist 13 when compared to vehicle (Figure 2A-G). This GPR4 antagonist 13-mediated reduction in histopathology occurred in both the distal and middle colon segments (P < 0.01) with a trend in reduction at the proximal segment, though not statically significant. Interestingly, the degree of leukocyte infiltration was also reduced by GPR4 antagonist 13 when compared to vehicle (Figure 2H).

**Figure 2:**
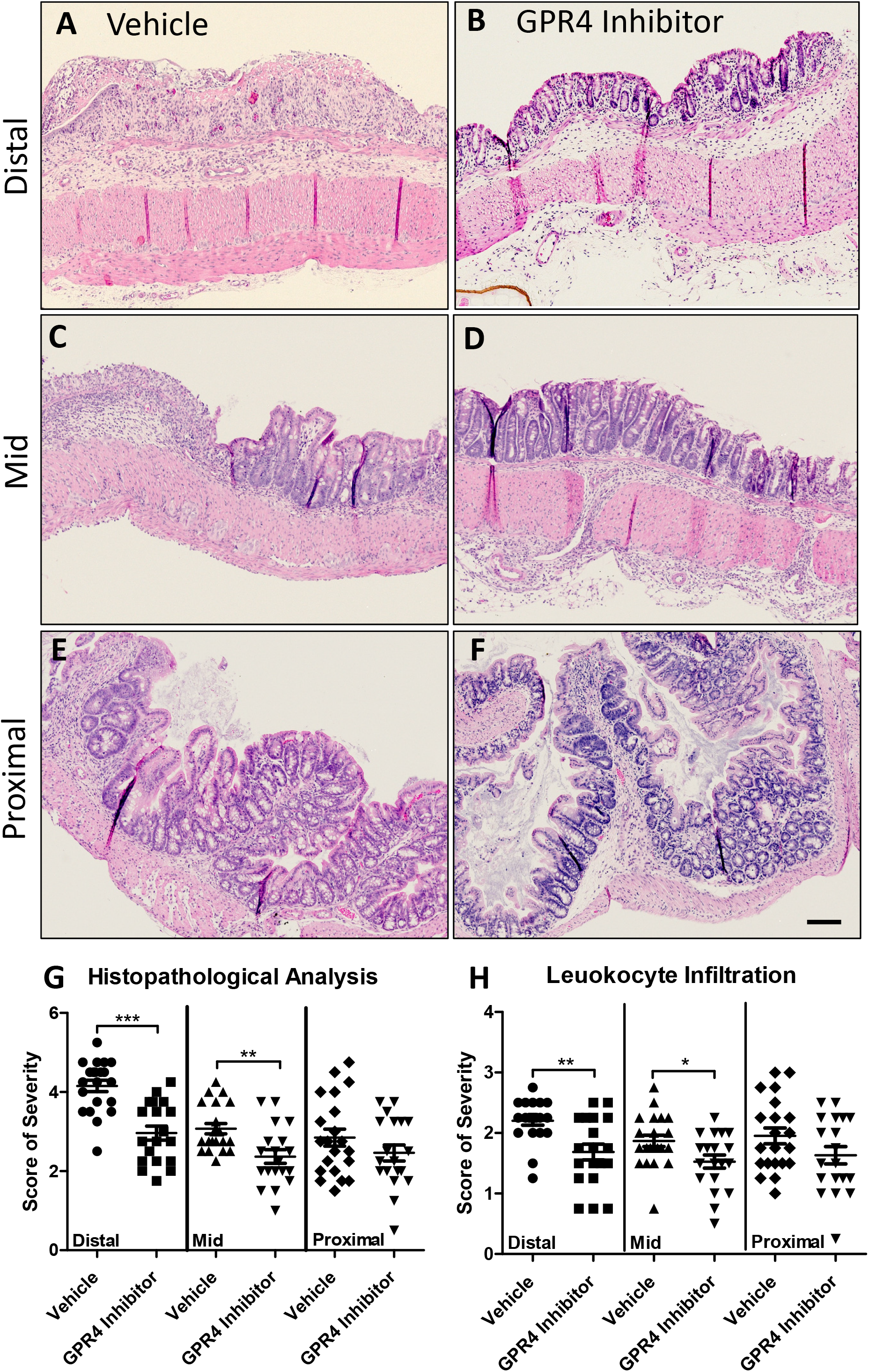
GPR4 antagonist 13 reduces histopathological parameters of intestinal inflammation in the inflamed mouse colon. Distinct histopathological features of intestinal inflammation were assessed and scored for degree of severity. Representative pictures of vehicle distal (A), Middle (C), and proximal (E) colon segments compared to GPR4 antagonist 13 distal (B), middle (D), and proximal (F) colon segments. Graphical representation of total histopathological parameters (G) and leukocyte infiltration score (H). Vehicle: N=21 (10 male/11 female) and GPR4 antagonist 13: N=19 (10male/9 female). Data are presented as mean ± SEM and was analyzed for statistical significance using the t-test between vehicle and GPR4 antagonist 13 groups between each colon segment. (*P < 0.05, **P < 0.01, *** P < 0.001). 10x objective. Scale bar = 100μm.

### 3.4. Endothelium specific VCAM-1 and E-selectin expression are reduced by GPR4 antagonist 13

Previous studies have demonstrated GPR4 activation in endothelial cells (ECs) induces vascular cell adhesion molecule-1 (VCAM-1) and E-selectin expression (Chen et al., 2011; Dong et al., 2013; Tobo et al., 2015). Additionally, reports have shown GPR4 genetic deletion can reduce VCAM-1 and E-selectin expression in the vascular endothelium (Sanderlin et al., 2017). Here we assessed the protein expression signal intensity of VCAM-1 and E-selectin of intestinal microvascular endothelial cells in the distal, middle, and proximal colon mucosa segments of vehicle and GPR4 antagonist 13 DSS-treated mice (Figures 3 and 4). The intensity scoring of VCAM-1 and E-selectin expression revealed highest signal within the distal intestinal endothelia with progressive intensity reduction from the middle and proximal colon, respectively. These data are consistent with the observation of reduced leukocyte infiltration in the middle and proximal colon segments compared to the distal segment (Figure 2H). Interestingly, VCAM-1 and E-selectin signal intensity in intestinal microvascular ECs was reduced in GPR4 antagonist 13 DSS-treated mice when compared to vehicle in the distal colon and a trend of reduction in the middle colon (Figures 3–4). Collectively, these data provide evidence GPR4 antagonist 13 reduces EC activation and subsequently leukocyte infiltration into the inflamed intestinal mucosa.

**Figure 3:**
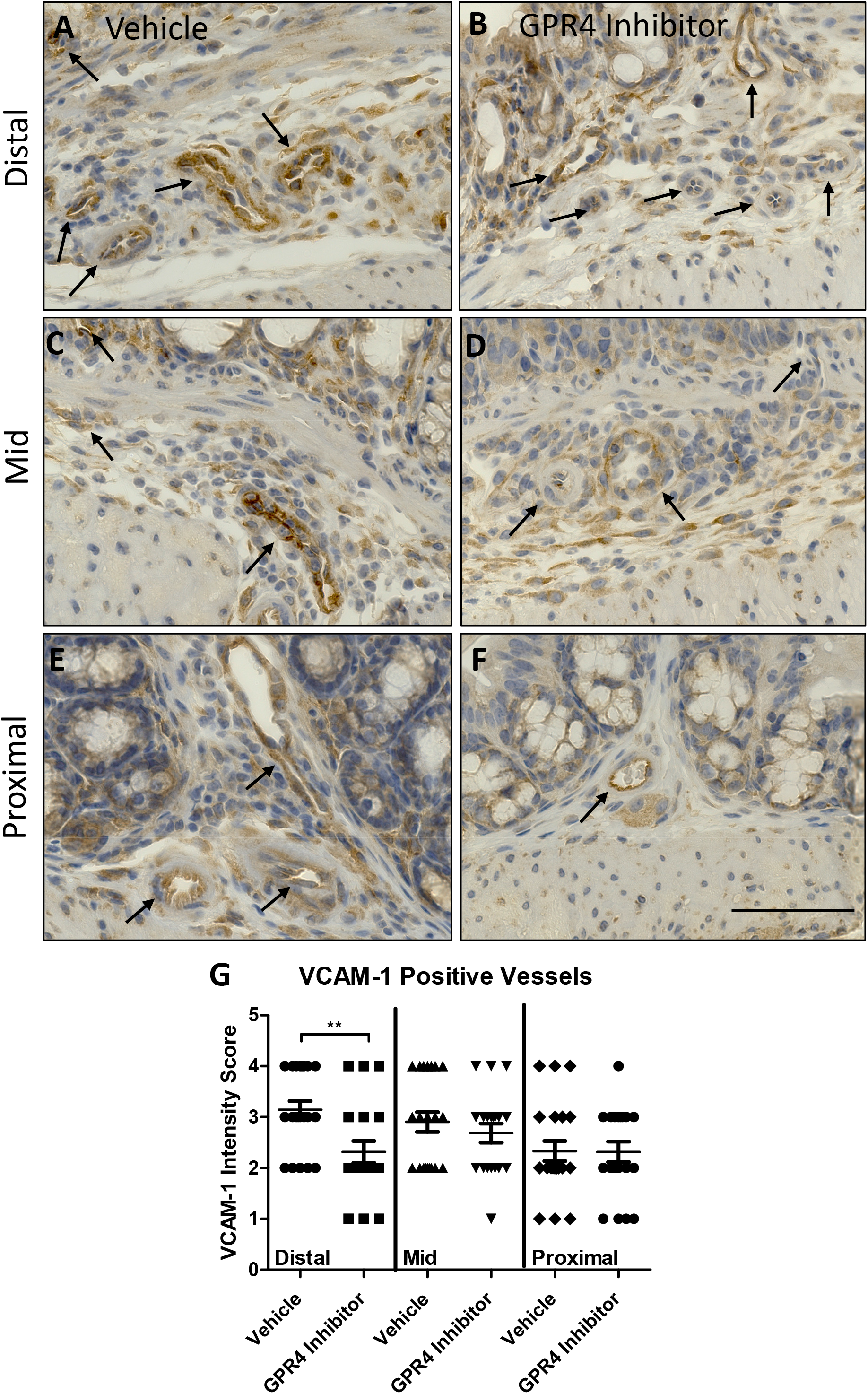
GPR4 antagonist 13 reduces VCAM-1 protein expression in colon microvascular endothelial cells. VCAM-1 protein expression intensity was assessed by immunohistochemistry (IHC) in colon microvascular endothelial cells. Representative pictures of vehicle distal (A), middle (C), and proximal (E) colon segments compared to GPR4 antagonist 13 distal (B), middle (D), and proximal (F) colon segments followed by graphical representation of VCAM-1 intensity score (G). Vehicle: N=21 (10 male/11 female) and GPR4 antagonist 13: N=19 (10male/9 female). Data are presented as mean ± SEM and was analyzed for statistical significance using the t-test between vehicle and GPR4 antagonist 13 groups between each colon segment. (**P < 0.01). 40x objective. Scale bar = 100μm. Black arrows indicate blood vessels.

**Figure 4:**
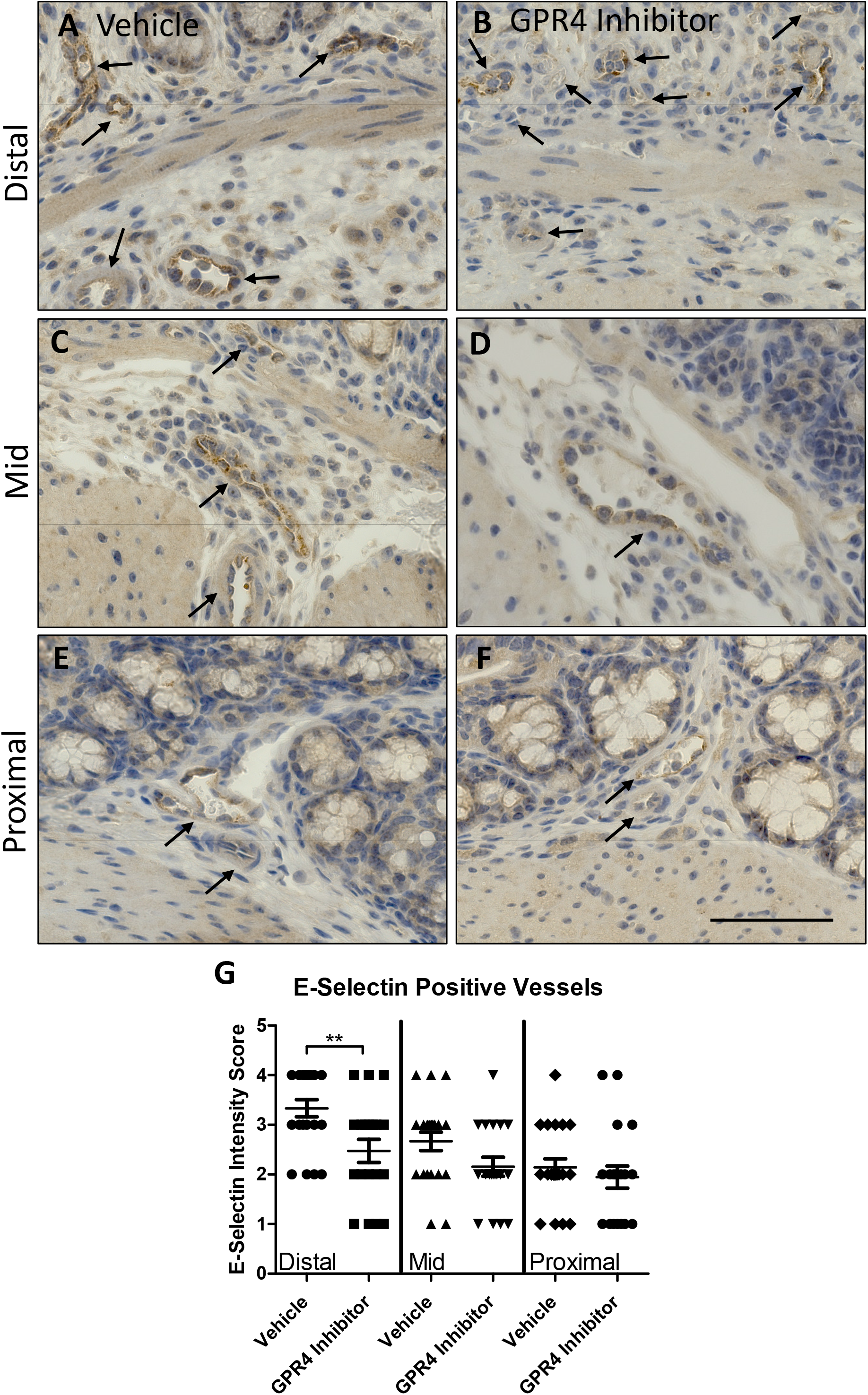
GPR4 antagonist 13 reduces E-selectin protein expression in colon microvascular endothelial cells. E-selectin protein expression intensity was assessed by IHC in colon microvascular endothelial cells. Representative pictures of vehicle distal (A), middle (C), and proximal (E) colon segments compared to GPR4 antagonist 13 distal (B), middle (D), and proximal (F) colon segments followed by graphical representation of E-selectin intensity score (G). Vehicle: N=21 (10 male/11 female) and GPR4 antagonist 13: N=19 (10male/9 female). Data are presented as mean ± SEM and was analyzed for statistical significance using the t-test between vehicle and GPR4 antagonist 13 groups between each colon segment. (**P < 0.01). 40x objective. Scale bar = 100μm. Black arrows indicate blood vessels.

Immunohistochemical analysis also revealed expression of VCAM-1 and E-selectin on cell types not regulated by GPR4. VCAM-1 expression could be strongly observed in activated fibroblasts and other mononuclear cells within the intestinal mucosa (Supplemental Figure 1A-B). Previous reports have observed expression of VCAM-1 in skeletal muscle, fibroblast, and some leukocyte populations (Epperly et al., 2002; Ulyanova et al., 2005). E-selectin expression could be detected within the colon epithelium (Supplemental Figure 1C-D). These results are consistent with previous studies showing E-selectin can be expressed on the colon epithelium and in mononuclear cells in the intestinal mucosa during active colitis (Vainer et al., 1998). Furthermore, additional studies have shown E-selectin is expressed in T cells and can be upregulated by pro-inflammatory mediators (Bajnok et al., 2017; Harashima et al., 2001). We observed similar levels of signal intensity of VCAM-1 and E-selectin in cell types other than intestinal microvascular endothelial cells between vehicle and GPR4 antagonist 13 groups (Supplemental Figure 1).

In addition to VCAM-1 and E-selectin expression analysis, we assessed the expression intensity and distribution of mucosal vascular addressin cell adhesion molecule-1 (MAdCAM-1) in colon tissues. We observed prominent MAdCAM-1 expression in intestinal microvasculature with high expression density in the lamina propria (Figure 5). No MAdCAM-1 expression could be observed in arteries and extramural blood vessels and minimal expression could be detected in lymphatic endothelial cells. MAdCAM-1 expression signal intensity in the intestinal microvasculature were similar between vehicle and GPR4 antagonist 13 DSS-treated mouse groups (Figure 5A-B). Total number of vessels positive for MAdCAM-1, however, were markedly reduced in GPR4 antagonist 13 mice compared to vehicle control mice (Figure 5A-D). Total MAdCAM-1 positive vessels were counted per centimeter (cm) from the entire length of the colon. Vehicle treated mice had ~60 MAdCAM-1 positive vessels/cm compared to ~35 MAdCAM-1 positive vessels/cm in GPR4 antagonist 13 treated mice (Figure 5E).

**Figure 5:**
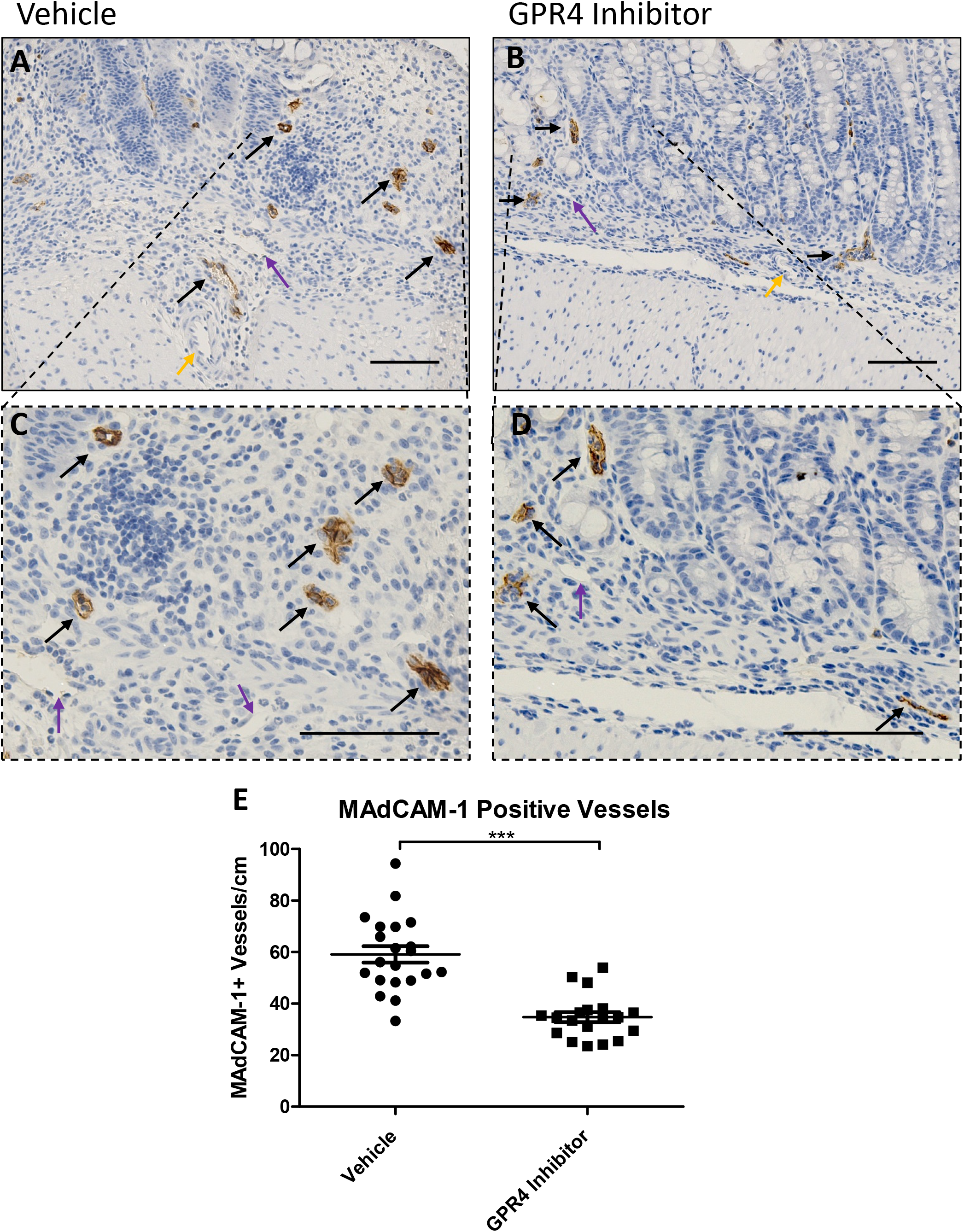
GPR4 antagonist 13 reduces MAdCAM-1 positive vessels in the mouse colon. No difference in MAdCAM-1 protein expression intensity was observed between vehicle and GPR4 antagonist 13 colon microvascular endothelial cells; however, differences in total number of MAdCAM-1 positive vessels were observed between the two treatment groups. Representative pictures of vehicle distal (A, C) compared to GPR4 antagonist 13 (B, D). Graphical representation of MAdCAM-1+ blood vessels per colon centimeter are depicted (E). Vehicle: N=21 (10 male/11 female) and GPR4 antagonist 13: N=19 (10male/9 female). Data are presented as mean ± SEM and was analyzed for statistical significance using the t-test between vehicle and GPR4 antagonist 13 groups. (*** P < 0.001). 20x and 40x objective. Black arrow indicates microvasculature, purple arrow indicates lymphatic endothelial cell, yellow arrow indicates artery. Scale bar = 100μm.

### 3.5. Inflammatory gene expression is reduced in the distal colon by GPR4 antagonist 13

Following the assessment of histopathology and endothelial cell-specific inflammatory protein expression, we assessed inflammatory gene expression at the whole tissue level to discern the anti-inflammatory effects of GPR4 antagonist 13. Inflammatory genes were measured from the distal colon segment of vehicle and GPR4 antagonist 13 DSS-treated mice. Additionally, distal colon segments of wild-type mice not treated with DSS were collected and assessed to serve as a baseline gene expression reference. Inflammatory mediators such as TNF-α, IL-1β, IL-6, IL-10, and PTGS2 (COX-2) were measured between vehicle and GPR4 antagonist 13 groups (Figure 6). A statistically significant reduction of TNF-α and IL-10 gene expression could be appreciated in the GPR4 antagonist 13 group compared to vehicle. Reduction of TNF-α is correlated with reduced inflammation in the mouse distal colon of the GPR4 antagonist 13 group. IL-10 can inhibit the function of macrophages and other inflammatory cells which are required for optimal pathogen clearance and subsequent inflammatory resolution (Couper et al., 2008; Kessler et al., 2017; Rojas et al., 2017). As such, reduced levels of IL-10 mRNA in the tissues of GPR4 antagonist 13 treated mice could potentially enhance pathogen clearance during the acute inflammatory phase. Furthermore, vehicle administered mice also had a trend in heighted expression of the inflammatory genes IL-1β and IL-6 when compared to the GPR4 antagonist 13 treated group. In addition to inflammatory cytokines, adhesion molecule gene expression including E-selectin, MAdCAM-1, and VCAM-1 were assessed. A statistically significant reduction of MAdCAM-1 expression could be discerned in the GPR4 antagonist 13 treated mice compared to vehicle (Figure 6B). These results are in line with reduced MAdCAM-1 positive vessels in the distal colon of GPR4 antagonist 13 mice compared to vehicle (Figure 5). VCAM-1 gene expression was modestly reduced in the GPR4 antagonist 13 group compared to vehicle and no trend in reduction could be discerned for E-selectin at the whole tissue level (Figure 6D, F). As VCAM-1 and E-selectin are expressed in cells that are not regulated by GPR4 (Supplemental Figure 1), whole tissue gene expression analysis presents complications for assessing VCAM-1 and E-selectin expression specific to GPR4 regulated vascular endothelial cells. To overcome this complication, we assessed VCAM-1 and E-selectin protein expression specific to vascular endothelial cells as described above (Figures 3–4). Taken together, the results indicate that GPR4 antagonist 13 reduces inflammatory gene expression in the distal colon of DSS-treated mice.

**Figure 6:**
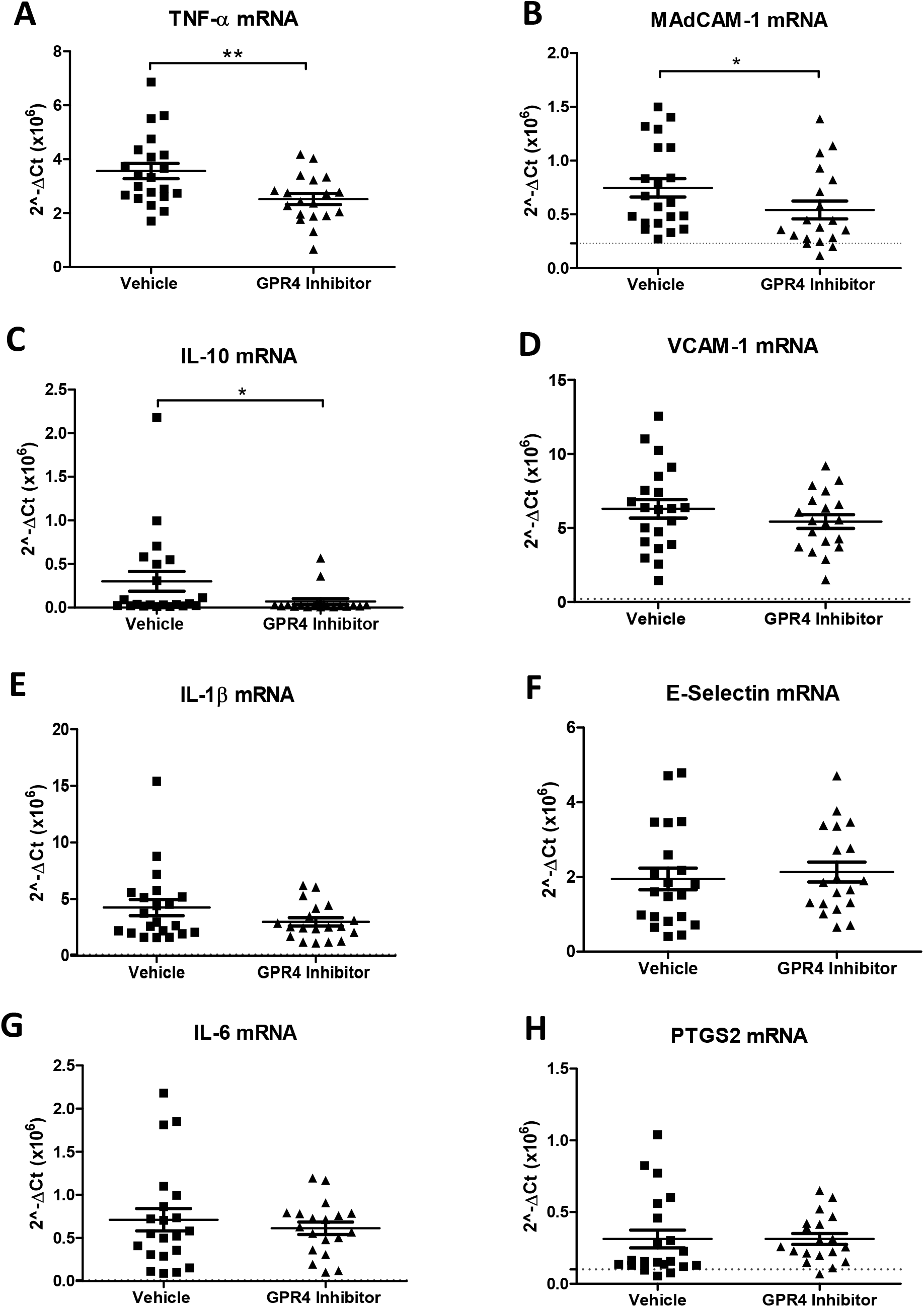
GPR4 antagonist 13 reduces inflammatory gene expression in the distal colon. Tissue level gene expression of cytokines, adhesion molecules, and an inflammatory enzyme were assessed in the distal colon segment of DSS-treated mice given vehicle or GPR4 antagonist 13. Graphical representation of TNF-α (A), MAdCAM-1 (B), IL-10 (C), VCAM-1 (D), IL-1β (E), E-selectin (F), IL-6 (G), and PTGS2 (H) gene expression. Vehicle: N=21 (10 male / 11 female) and GPR4 antagonist 13: N=19 (10male / 9 female). Data are presented as mean ± SEM and was analyzed for statistical significance using the Mann-Whitney test between vehicle and GPR4 antagonist 13 groups between each colon segment. (*P < 0.05, **P < 0.01). Dotted black line are indicative of untreated control C57Bl/6 mice (N= 2 male/2 female).

## 4. Discussion

The local pH of the inflammatory tissue is characteristically acidic (Lardner, 2001; Okajima, 2013), and can fluctuate within the physiological range to severely acidic. Some groups have evaluated the pH within gastrointestinal tract of patients with active IBD (Fallingborg et al., 1993; Nugent et al., 2001). These reports suggest the intraluminal colonic pH can be acidic in patients with active colitis. Molecular mechanisms for sensing extracellular pH during intestinal inflammatory responses have only recently been investigated.

The pH-sensing GPCRs, including GPR65 (TDAG8), GPR68 (OGR1), and GPR4, have emerged as regulators of intestinal inflammation (de Valliere et al., 2015; Hutter et al., 2018; Lassen et al., 2016; Sanderlin et al., 2017; Wang et al., 2018). Initial studies on the effects of acidic pH-induced activation of GPR65 have provided an anti-inflammatory role for GPR65 in macrophages and other immune cells (Jin et al., 2014; Mogi et al., 2009). Furthermore, genetic variances within the GPR65 gene were correlated to increased risk of the development of a variety of inflammatory diseases including inflammatory bowel disease (Franke et al., 2010; Jostins et al., 2012; Lassen et al., 2016). These data implicate pH homeostasis and cellular pH sensing mechanisms in the regulation of inflammation. A recent study investigated GPR65 in a bacteria-induced colitis model and found GPR65 protects against intestinal inflammation (Lassen et al., 2016). GPR68 (OGR1) has also been investigated and found to potentiate intestinal inflammation through regulating macrophage inflammatory programs in response to acidic pH (de Valliere et al., 2015). Finally, GPR4 has been studied in both acute and chronic intestinal inflammation mouse models (Sanderlin et al., 2017; Wang et al., 2018). Previous reports indicate GPR4 is predominately expressed in vascular endothelial cells (ECs) (Chen et al., 2011; Dong et al., 2013; Yang et al., 2007). GPR4 expression was evaluated within normal and inflamed mouse colon tissues and was found to be predominately expressed within intestinal ECs (Sanderlin et al., 2017). Several studies implicate GPR4 as a regulator of EC activation in response to extracellular acidosis by increasing vascular adhesion molecules such as VCAM-1, E-selectin, and ICAM-1 and functionally increasing EC-leukocyte interactions (Chen et al., 2011; Dong et al., 2013; Tobo et al., 2015). Previous studies have implicated these mechanistic insights into mouse colitis models. We have previously demonstrated genetic deletion of GPR4 in mice reduced intestinal inflammation in an acute DSS-induced colitis mouse model (Sanderlin et al., 2017). Additionally, GPR4 knockout (KO) mice had reduced neutrophil, macrophage, and T cell infiltration into the intestinal mucosa. The reduced leukocyte infiltration was associated with GPR4-dependent reduction of vascular adhesion molecule expression in the intestinal ECs. Another group assessed the functional role of GPR4 in chronic DSS-induced inflammation and the IL-10 KO spontaneous colitis mouse model (Wang et al., 2018). Mice devoid of GPR4 had reduced intestinal inflammation and subsequent mucosal CD4+ T helper cell infiltration. These studies collectively suggest GPR4 mediates intestinal inflammation through increasing gut leukocyte recruitment and subsequent extravasation into the intestinal mucosa. Inhibition of GPR4 may be explored as a novel anti-inflammatory approach.

In this study we show that the GPR4 antagonist 13 can reduce intestinal inflammation in the DSS-induced acute colitis mouse model. Upon initiation of DSS-induced colitis, mice receiving the GPR4 antagonist were protected from body weight loss, fecal blood and diarrhea, colon shortening, mesenteric lymph node expansion, and splenic enlargement. Colon histopathology was also alleviated. Moreover, a reduction of VCAM-1 and E-selectin expression in intestinal ECs as well as a reduced number of MAdCAM-1 positive microvessels were observed within the distal colon of mice given the GPR4 antagonist. These observations suggest that the GPR4 antagonist 13 reduces the expression of vascular adhesion molecules on ECs and thus attenuates leukocyte-endothelium adhesion and extravasation (Figure 7).

**Figure 7:**
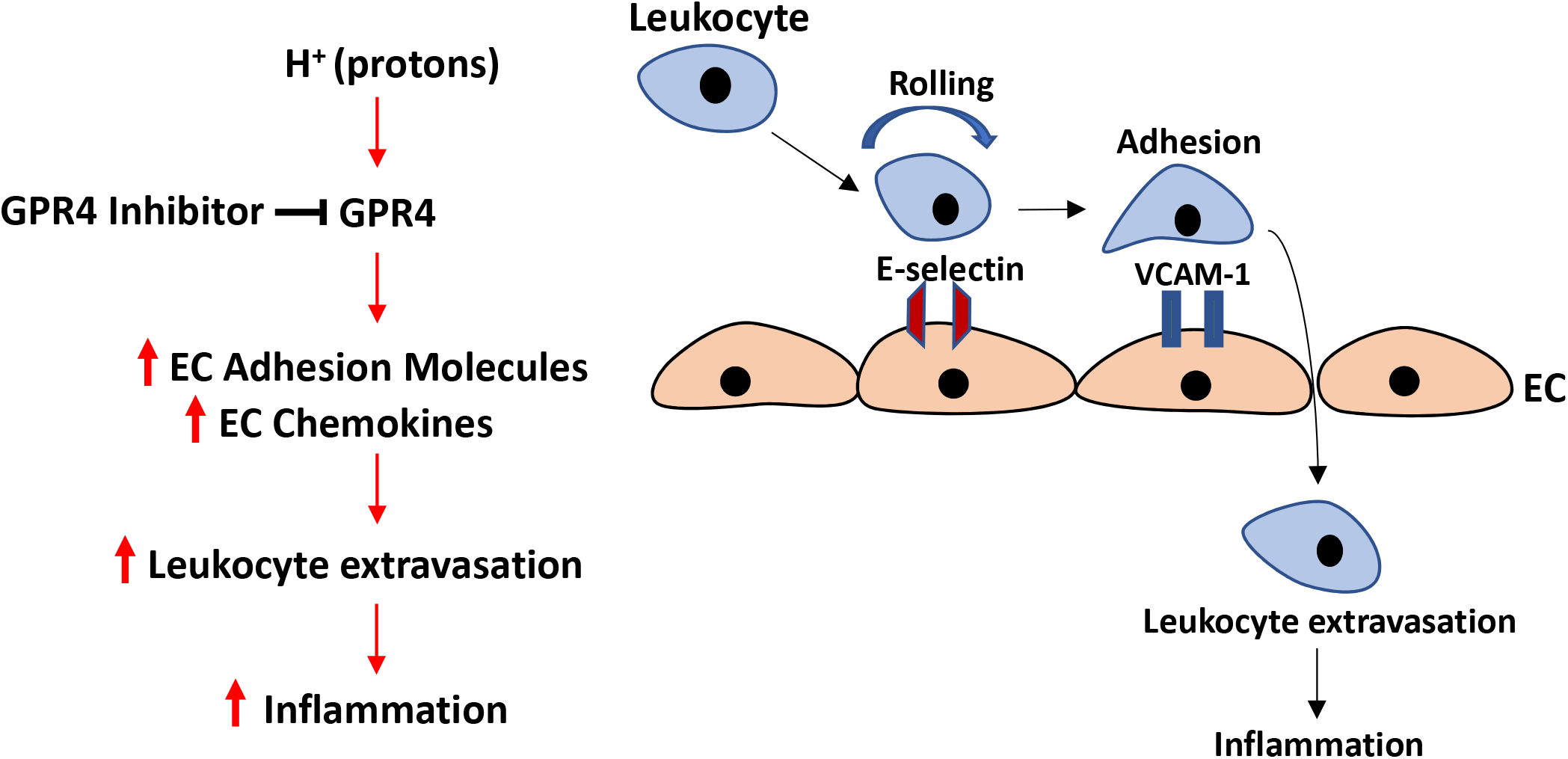
Model of proposed mechanism of the anti-inflammatory action of GPR4 inhibitors. GPR4 activation by protons in the extracellular milieu mediates the activation of vascular endothelial cells, the recruitment of immune cells and subsequent leukocyte extravasation into the inflamed tissue. Heavy immune cell infiltration into the inflammatory loci will result in further production of protons, as well as pro-inflammatory mediators, and subsequently maintain tissue inflammation and GPR4 activation. Inhibition of GPR4 activity by pharmacological intervention may present a novel approach to reduce inflammation by attenuating vascular endothelial cell activation and leukocyte infiltration into inflamed tissues. EC: endothelial cell.

Upon immunohistochemical evaluation of VCAM-1 and E-selectin within the colon, notable expression could also be detected in cell types not known to be regulated by GPR4. VCAM-1 could be observed on fibroblasts, macrophages, and other mononuclear cells within the colon. These observations are supported by several other studies describing basal and inflammation-associated VCAM-1 upregulation on activated fibroblasts and inflammatory cells (Epperly et al., 2002; Ulyanova et al., 2005). Studies have also observed that E-selectin can be expressed by the colonic epithelium and mononuclear cells including T cells (Bajnok et al., 2017; Harashima et al., 2001; Vainer et al., 1998). Our results were consistent with these reports as E-selectin could be detected in colonic epithelial cells and mononuclear cell populations within the inflamed mucosa (Supplemental Figure 1). Notably, the expression of VCAM-1 and E-selectin in fibroblasts, colonic epithelial cells and mononuclear cells was at a similar level in the colon of GPR4 antagonist 13 and vehicle treated mice. These observations may explain why the mRNA expression levels of VCAM-1 and E-selectin in the whole colon tissue were either just slightly reduced or not changed in the mice treated with GPR4 antagonist 13 compared to vehicle as these cell types are not regulated by GPR4.

In addition to vascular adhesion molecules, the expression of TNF-α and IL-10 was also significantly reduced in the distal colon tissue of mice given the GPR4 antagonist. TNF-α and IL-10 mRNA reduction at the whole tissue level is most likely due to the reduction of inflammatory cell infiltration into the colon observed in the GPR4 antagonist 13 treated mice. Increased IL-10 mRNA levels are commonly observed in the acute DSS-induced colitis mouse model and in mucosal T cells of patients with active ulcerative colitis (Egger et al., 2000; Melgar et al., 2003). GPR4 antagonist 13-mediated reduction of IL-10 mRNA is in line with reduced leukocyte infiltration. Furthermore, as mentioned above, reduced IL-10 mRNA levels could aid optimal pathogen clearance during the acute inflammatory phase (Couper et al., 2008; Kessler et al., 2017; Rojas et al., 2017). A modest reduction of IL-1β and IL-6 could also be observed in the colon of mice treated with the GPR4 antagonist. Collectively, these results implicate GPR4 inhibitors as novel anti-inflammatory agents for colitis-associated intestinal inflammation.

GPR4 inhibitors have proved effective in the reduction of endothelial inflammation and subsequent tissue inflammation *in vitro* and *in vivo* (Dong et al., 2017; Dong et al., 2013; Fukuda et al., 2016; Hosford et al., 2018; Miltz et al., 2017; Tobo et al., 2015; Velcicky et al., 2017). A group of imidazo-pyridine and pyrazolopyrimidine derivatives were previously found to elicit inhibitory effects on GPR4-mediated EC activation (Dong et al., 2017; Dong et al., 2013; Hosford et al., 2018; Miltz et al., 2017; Tobo et al., 2015; Velcicky et al., 2017). Further studies revealed this class of compounds could reduce mortality events in a myocardial infraction mouse model (Fukuda et al., 2016). Recently, the imidazo-pyridine derivatives were further characterized as an orally active GPR4 antagonist 39c (Miltz et al., 2017). This compound is commercially available and used to provide proof of concept in the pro-inflammatory role of GPR4 in endothelial inflammation (Dong et al., 2017; Dong et al., 2013). However, due to off-target effects such as cross-reactivity with the histamine H3 receptor, GPR4 antagonist 39c was not further pursued (Miltz et al., 2017; Velcicky et al., 2017). Finally, the GPR4 antagonist 13, a pyrazolopyrimidine derivative, was developed and evaluated (Velcicky et al., 2017). Common off-targets, such as the H3 receptor and hERG channel as well as other pH-sensing GPCRs, were evaluated for GPR4 antagonist 13 and the compound proved selective to the GPR4 receptor (Velcicky et al., 2017). *In vivo* pharmacokinetic studies demonstrated GPR4 antagonist 13 had good profiles of oral delivery and drug clearance. Additionally, an *in vitro* safety profile assessment against a panel of receptors, enzymes, and ion channels indicated GPR4 antagonist 13 did not significantly inhibit these targets. Furthermore, GPR4 antagonist 13 was found to inhibit arthritis inflammation as well as hyperplasia and angiogenesis in animal models (Velcicky et al., 2017). Taken together, the optimized GPR4 antagonist 13 displays a good pharmacological profile and has proven orally active for *in vivo* use. Nonetheless, there are some aspects requiring close attention for the use of GPR4 inhibitors.

Previous studies have shown that a fraction of GPR4 knockout mice display vascular abnormalities associated with perinatal complications (Yang et al., 2007). GPR4-null mice also exhibit decreased renal acid excretion (Sun et al., 2010). Other reports have shown GPR4 is involved in CO_2_ chemosensing to regulate breathing (Kumar et al., 2015). A recent study evaluated the central nervous system (CNS) effects of GPR4 antagonist 13 and observed no effects on hemodynamics, blood oxygen levels, and cerebral blood flow (Hosford et al., 2018). However, GPR4 antagonist 13 reduced hypercapnic response to CO_2_ but had no effect when mice were under anesthesia. These studies suggest that current GPR4 inhibitors have a good overall pharmacological profile but might not be taken during pregnancy or if renal and respiratory complications exist.

In summary, this study is the first to evaluate the effects of the pharmacological inhibition of GPR4 on intestinal inflammation. We have shown that GPR4 antagonization can reduce the degree of intestinal inflammation in the DSS-induced acute colitis mouse model.

## Acknowledgements

This work was supported by a grant from the National Institutes of Health (R15DK109484, to L.V.Y).

## Author contributions

E.J.S. and L.V.Y. designed the experiments; E.J.S. and M.M. performed the experiments; E.J.S., M.M. and L.V.Y. analyzed the data; J.V. and P.L. provided the GPR4 antagonist 13 and instructions on how to administer the antagonist in mice; E.J.S. and L.V.Y. wrote the manuscript; all authors reviewed and approved the manuscript.

---

**Supplemental Figure 1:**
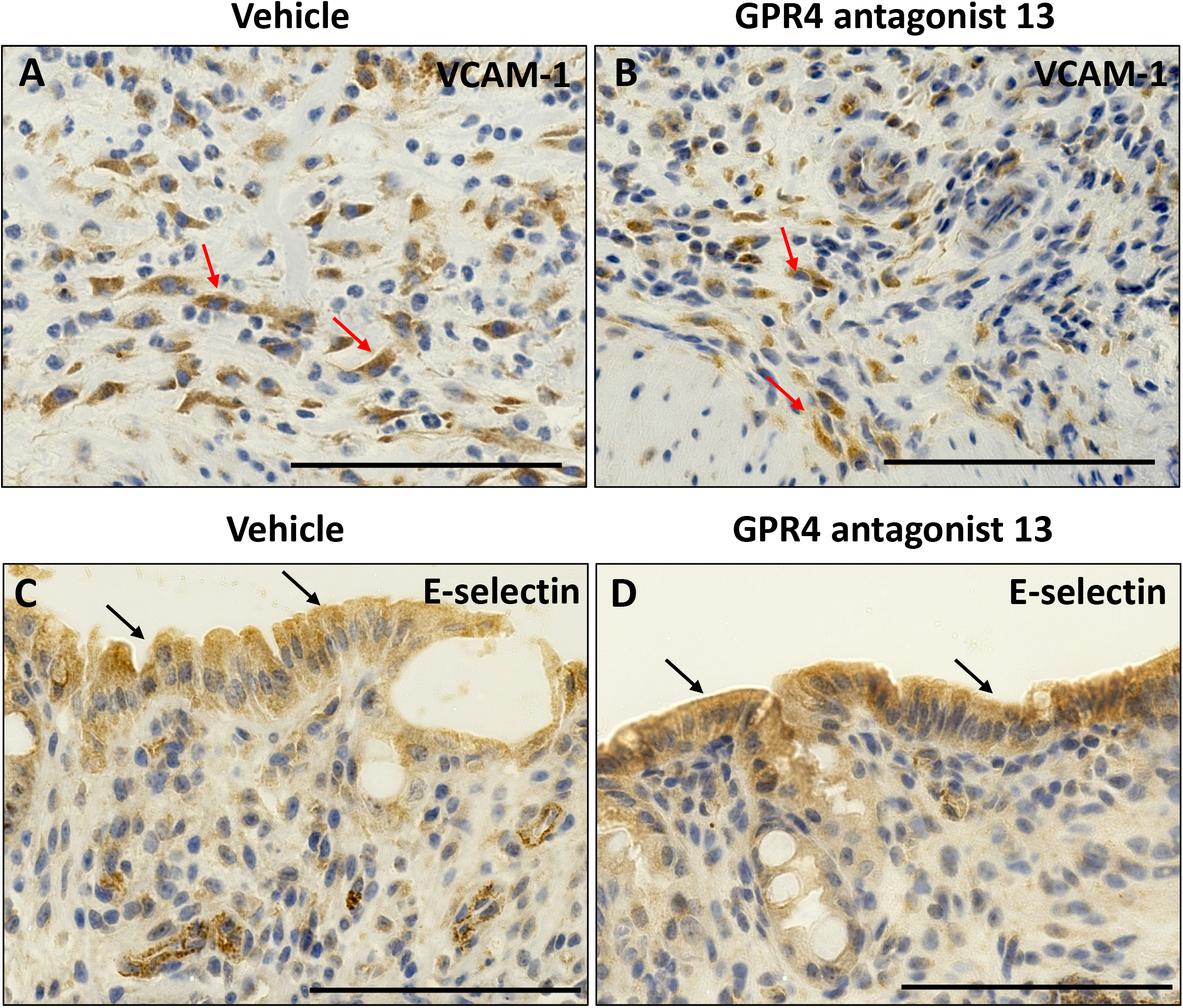
GPR4 antagonist 13 does not alter VCAM-1 or E-selectin expression in cell types that are not known to be regulated by GPR4. VCAM-1 and E-selectin expression was assessed in non-endothelial cells within the colon. VCAM-1 is expressed in non-endothelial mononuclear cells and fibroblasts and E-selectin is expressed in the intestinal epithelium. Representative pictures of VCAM-1 in vehicle (A) and GPR4 antagonist 13-treated mice (B) and E-selectin in vehicle (C) and GPR4 antagonist 13-treated mice (D) in the colon. Scale bar = 100μm. Red arrows indicate VCAM-1 positive cells and black arrows indicate E-selectin positive cells.

## References

Bajnok, A., Ivanova, M., Rigo, J., Jr., Toldi, G., 2017. The Distribution of Activation Markers and Selectins on Peripheral T Lymphocytes in Preeclampsia. Mediators Inflamm 2017, 8045161.

Chassaing, B., Aitken, J.D., Malleshappa, M., Vijay-Kumar, M., 2014. Dextran sulfate sodium (DSS)-induced colitis in mice. Curr Protoc Immunol 104, Unit 15 25.

Chen, A., Dong, L., Leffler, N.R., Asch, A.S., Witte, O.N., Yang, L.V., 2011. Activation of GPR4 by acidosis increases endothelial cell adhesion through the cAMP/Epac pathway. PLoS One 6, e27586.

Couper, K.N., Blount, D.G., Riley, E.M., 2008. IL-10: the master regulator of immunity to infection. J Immunol 180, 5771–5777.

de Valliere, C., Wang, Y., Eloranta, J.J., Vidal, S., Clay, I., Spalinger, M.R., Tcymbarevich, I., Terhalle, A., Ludwig, M.G., Suply, T., Fried, M., Kullak-Ublick, G.A., Frey-Wagner, I., Scharl, M., Seuwen, K., Wagner, C.A., Rogler, G., 2015. G Protein-coupled pH-sensing Receptor OGR1 Is a Regulator of Intestinal Inflammation. Inflamm Bowel Dis 21, 1269–1281.

Dong, L., Krewson, E.A., Yang, L.V., 2017. Acidosis Activates Endoplasmic Reticulum Stress Pathways through GPR4 in Human Vascular Endothelial Cells. Int J Mol Sci 18.

Dong, L., Li, Z., Leffler, N.R., Asch, A.S., Chi, J.T., Yang, L.V., 2013. Acidosis Activation of the Proton-Sensing GPR4 Receptor Stimulates Vascular Endothelial Cell Inflammatory Responses Revealed by Transcriptome Analysis. PLoS One 8, e61991.

Egger, B., Bajaj-Elliott, M., MacDonald, T.T., Inglin, R., Eysselein, V.E., Buchler, M.W., 2000. Characterisation of acute murine dextran sodium sulphate colitis: cytokine profile and dose dependency. Digestion 62, 240–248.

Epperly, M.W., Sikora, C.A., DeFilippi, S.J., Gretton, J.E., Bar-Sagi, D., Archer, H., Carlos, T., Guo, H., Greenberger, J.S., 2002. Pulmonary irradiation-induced expression of VCAM-I and ICAM-I is decreased by manganese superoxide dismutase-plasmid/liposome (MnSOD-PL) gene therapy. Biol Blood Marrow Transplant 8, 175–187.

Erben, U., Loddenkemper, C., Doerfel, K., Spieckermann, S., Haller, D., Heimesaat, M.M., Zeitz, M., Siegmund, B., Kuhl, A.A., 2014. A guide to histomorphological evaluation of intestinal inflammation in mouse models. Int J Clin Exp Pathol 7, 4557–4576.

Fallingborg, J., Christensen, L.A., Jacobsen, B.A., Rasmussen, S.N., 1993. Very low intraluminal colonic pH in patients with active ulcerative colitis. Dig Dis Sci 38, 1989–1993.

Franke, A., McGovern, D.P., Barrett, J.C., Wang, K., Radford-Smith, G.L., Ahmad, T., Lees, C.W., Balschun, T., Lee, J., Roberts, R., Anderson, C.A., Bis, J.C., Bumpstead, S., Ellinghaus, D., Festen, E.M., Georges, M., Green, T., Haritunians, T., Jostins, L., Latiano, A., Mathew, C.G., Montgomery, G.W., Prescott, N.J., Raychaudhuri, S., Rotter, J.I., Schumm, P., Sharma, Y., Simms, L.A., Taylor, K.D., Whiteman, D., Wijmenga, C., Baldassano, R.N., Barclay, M., Bayless, T.M., Brand, S., Buning, C., Cohen, A., Colombel, J.F., Cottone, M., Stronati, L., Denson, T., De Vos, M., D’Inca, R., Dubinsky, M., Edwards, C., Florin, T., Franchimont, D., Gearry, R., Glas, J., Van Gossum, A., Guthery, S.L., Halfvarson, J., Verspaget, H.W., Hugot, J.P., Karban, A., Laukens, D., Lawrance, I., Lemann, M., Levine, A., Libioulle, C., Louis, E., Mowat, C., Newman, W., Panes, J., Phillips, A., Proctor, D.D., Regueiro, M., Russell, R., Rutgeerts, P., Sanderson, J., Sans, M., Seibold, F., Steinhart, A.H., Stokkers, P.C., Torkvist, L., Kullak-Ublick, G., Wilson, D., Walters, T., Targan, S.R., Brant, S.R., Rioux, J.D., D’Amato, M., Weersma, R.K., Kugathasan, S., Griffiths, A.M., Mansfield, J.C., Vermeire, S., Duerr, R.H., Silverberg, M.S., Satsangi, J., Schreiber, S., Cho, J.H., Annese, V., Hakonarson, H., Daly, M.J., Parkes, M., 2010. Genome-wide meta-analysis increases to 71 the number of confirmed Crohn’s disease susceptibility loci. Nat Genet 42, 1118–1125.

Fukuda, H., Ito, S., Watari, K., Mogi, C., Arisawa, M., Okajima, F., Kurose, H., Shuto, S., 2016. Identification of a Potent and Selective GPR4 Antagonist as a Drug Lead for the Treatment of Myocardial Infarction. ACS Med Chem Lett 7, 493–497.

Harashima, S., Horiuchi, T., Hatta, N., Morita, C., Higuchi, M., Sawabe, T., Tsukamoto, H., Tahira, T., Hayashi, K., Fujita, S., Niho, Y., 2001. Outside-to-inside signal through the membrane TNF-alpha induces E-selectin (CD62E) expression on activated human CD4+ T cells. J Immunol 166, 130–136.

Hendrickson, B.A., Gokhale, R., Cho, J.H., 2002. Clinical aspects and pathophysiology of inflammatory bowel disease. Clin Microbiol Rev 15, 79–94.

Hosford, P.S., Mosienko, V., Kishi, K., Jurisic, G., Seuwen, K., Kinzel, B., Ludwig, M.G., Wells, J.A., Christie, I.N., Koolen, L., Abdala, A.P., Liu, B.H., Gourine, A.V., Teschemacher, A.G., Kasparov, S., 2018. CNS distribution, signalling properties and central effects of G-protein coupled receptor 4. Neuropharmacology 138, 381–392.

Hutter, S., van Haaften, W.T., Hunerwadel, A., Baebler, K., Herfarth, N., Raselli, T., Mamie, C., Misselwitz, B., Rogler, G., Weder, B., Dijkstra, G., Meier, C.F., de Valliere, C., Weber, A., Imenez Silva, P.H., Wagner, C.A., Frey-Wagner, I., Ruiz, P.A., Hausmann, M., 2018. Intestinal activation of pH-sensing receptor OGR1 (GPR68) contributes to fibrogenesis. J Crohns Colitis.

Jin, Y., Sato, K., Tobo, A., Mogi, C., Tobo, M., Murata, N., Ishii, S., Im, D.S., Okajima, F., 2014. Inhibition of interleukin-1beta production by extracellular acidification through the TDAG8/cAMP pathway in mouse microglia. J Neurochem 129, 683–695.

Jostins, L., Ripke, S., Weersma, R.K., Duerr, R.H., McGovern, D.P., Hui, K.Y., Lee, J.C., Schumm, L.P., Sharma, Y., Anderson, C.A., Essers, J., Mitrovic, M., Ning, K., Cleynen, I., Theatre, E., Spain, S.L., Raychaudhuri, S., Goyette, P., Wei, Z., Abraham, C., Achkar, J.P., Ahmad, T., Amininejad, L., Ananthakrishnan, A.N., Andersen, V., Andrews, J.M., Baidoo, L., Balschun, T., Bampton, P.A., Bitton, A., Boucher, G., Brand, S., Buning, C., Cohain, A., Cichon, S., D’Amato, M., De Jong, D., Devaney, K.L., Dubinsky, M., Edwards, C., Ellinghaus, D., Ferguson, L.R., Franchimont, D., Fransen, K., Gearry, R., Georges, M., Gieger, C., Glas, J., Haritunians, T., Hart, A., Hawkey, C., Hedl, M., Hu, X., Karlsen, T.H., Kupcinskas, L., Kugathasan, S., Latiano, A., Laukens, D., Lawrance, I.C., Lees, C.W., Louis, E., Mahy, G., Mansfield, J., Morgan, A.R., Mowat, C., Newman, W., Palmieri, O., Ponsioen, C.Y., Potocnik, U., Prescott, N.J., Regueiro, M., Rotter, J.I., Russell, R.K., Sanderson, J.D., Sans, M., Satsangi, J., Schreiber, S., Simms, L.A., Sventoraityte, J., Targan, S.R., Taylor, K.D., Tremelling, M., Verspaget, H.W., De Vos, M., Wijmenga, C., Wilson, D.C., Winkelmann, J., Xavier, R.J., Zeissig, S., Zhang, B., Zhang, C.K., Zhao, H., Silverberg, M.S., Annese, V., Hakonarson, H., Brant, S.R., Radford-Smith, G., Mathew, C.G., Rioux, J.D., Schadt, E.E., Daly, M.J., Franke, A., Parkes, M., Vermeire, S., Barrett, J.C., Cho, J.H., 2012. Host-microbe interactions have shaped the genetic architecture of inflammatory bowel disease. Nature 491, 119–124.

Justus, C.R., Dong, L., Yang, L.V., 2013. Acidic tumor microenvironment and pH-sensing G protein-coupled receptors. Front Physiol 4, 354.

Justus, C.R., Sanderlin, E.J., Dong, L., Sun, T., Chi, J.T., Lertpiriyapong, K., Yang, L.V., 2017. Contextual tumor suppressor function of T cell death-associated gene 8 (TDAG8) in hematological malignancies. J Transl Med 15, 204.

Justus, C.R., Sanderlin, E.J., Yang, L.V., 2015. Molecular Connections between Cancer Cell Metabolism and the Tumor Microenvironment. Int J Mol Sci 16, 11055–11086.

Kaser, A., Zeissig, S., Blumberg, R.S., 2010. Inflammatory bowel disease. Annu Rev Immunol 28, 573–621.

Kessler, B., Rinchai, D., Kewcharoenwong, C., Nithichanon, A., Biggart, R., Hawrylowicz, C.M., Bancroft, G.J., Lertmemongkolchai, G., 2017. Interleukin 10 inhibits pro-inflammatory cytokine responses and killing of Burkholderia pseudomallei. Sci Rep 7, 42791.

Kim, J.J., Shajib, M.S., Manocha, M.M., Khan, W.I., 2012. Investigating intestinal inflammation in DSS-induced model of IBD. J Vis Exp.

Kumar, N.N., Velic, A., Soliz, J., Shi, Y., Li, K., Wang, S., Weaver, J.L., Sen, J., Abbott, S.B., Lazarenko, R.M., Ludwig, M.G., Perez-Reyes, E., Mohebbi, N., Bettoni, C., Gassmann, M., Suply, T., Seuwen, K., Guyenet, P.G., Wagner, C.A., Bayliss, D.A., 2015. PHYSIOLOGY. Regulation of breathing by CO(2) requires the proton-activated receptor GPR4 in retrotrapezoid nucleus neurons. Science 348, 1255–1260.

Lardner, A., 2001. The effects of extracellular pH on immune function. J Leukoc Biol 69, 522–530.

Laroui, H., Ingersoll, S.A., Liu, H.C., Baker, M.T., Ayyadurai, S., Charania, M.A., Laroui, F., Yan, Y., Sitaraman, S.V., Merlin, D., 2012. Dextran sodium sulfate (DSS) induces colitis in mice by forming nano-lipocomplexes with medium-chain-length fatty acids in the colon. PLoS One 7, e32084.

Lassen, K.G., McKenzie, C.I., Mari, M., Murano, T., Begun, J., Baxt, L.A., Goel, G., Villablanca, E.J., Kuo, S.Y., Huang, H., Macia, L., Bhan, A.K., Batten, M., Daly, M.J., Reggiori, F., Mackay, C.R., Xavier, R.J., 2016. Genetic Coding Variant in GPR65 Alters Lysosomal pH and Links Lysosomal Dysfunction with Colitis Risk. Immunity.

Ludwig, M.G., Vanek, M., Guerini, D., Gasser, J.A., Jones, C.E., Junker, U., Hofstetter, H., Wolf, R.M., Seuwen, K., 2003. Proton-sensing G-protein-coupled receptors. Nature 425, 93–98.

Mattar, M.C., Lough, D., Pishvaian, M.J., Charabaty, A., 2011. Current management of inflammatory bowel disease and colorectal cancer. Gastrointest Cancer Res 4, 53–61.

Melgar, S., Yeung, M.M., Bas, A., Forsberg, G., Suhr, O., Oberg, A., Hammarstrom, S., Danielsson, A., Hammarstrom, M.L., 2003. Over-expression of interleukin 10 in mucosal T cells of patients with active ulcerative colitis. Clin Exp Immunol 134, 127–137.

Miltz, W., Velcicky, J., Dawson, J., Littlewood-Evans, A., Ludwig, M.G., Seuwen, K., Feifel, R., Oberhauser, B., Meyer, A., Gabriel, D., Nash, M., Loetscher, P., 2017. Design and synthesis of potent and orally active GPR4 antagonists with modulatory effects on nociception, inflammation, and angiogenesis. Bioorg Med Chem 25, 4512–4525.

Mogi, C., Tobo, M., Tomura, H., Murata, N., He, X.D., Sato, K., Kimura, T., Ishizuka, T., Sasaki, T., Sato, T., Kihara, Y., Ishii, S., Harada, A., Okajima, F., 2009. Involvement of proton-sensing TDAG8 in extracellular acidification-induced inhibition of proinflammatory cytokine production in peritoneal macrophages. J Immunol 182, 3243–3251.

Neurath, M.F., 2017. Current and emerging therapeutic targets for IBD. Nat Rev Gastroenterol Hepatol 14, 269–278.

Nugent, S.G., Kumar, D., Rampton, D.S., Evans, D.F., 2001. Intestinal luminal pH in inflammatory bowel disease: possible determinants and implications for therapy with aminosalicylates and other drugs. Gut 48, 571–577.

Okajima, F., 2013. Regulation of inflammation by extracellular acidification and proton-sensing GPCRs. Cell Signal 25, 2263–2271.

Perse, M., Cerar, A., 2012. Dextran sodium sulphate colitis mouse model: traps and tricks. J Biomed Biotechnol 2012, 718617.

Rojas, J.M., Avia, M., Martin, V., Sevilla, N., 2017. IL-10: A Multifunctional Cytokine in Viral Infections. J Immunol Res 2017, 6104054.

Sanderlin, E.J., Justus, C.R., Krewson, E.A., Yang, L.V., 2015. Emerging roles for the pH-sensing G protein-coupled receptors in response to acidotic stress. Cell Health Cytoskelet 7, 99–109.

Sanderlin, E.J., Leffler, N.R., Lertpiriyapong, K., Cai, Q., Hong, H., Bakthavatchalu, V., Fox, J.G., Oswald, J.Z., Justus, C.R., Krewson, E.A., O’Rourke, D., Yang, L.V., 2017. GPR4 deficiency alleviates intestinal inflammation in a mouse model of acute experimental colitis. Biochim Biophys Acta 1863, 569–584.

Sun, X., Yang, L.V., Tiegs, B.C., Arend, L.J., McGraw, D.W., Penn, R.B., Petrovic, S., 2010. Deletion of the pH sensor GPR4 decreases renal acid excretion. J Am Soc Nephrol 21, 1745–1755.

Tobo, A., Tobo, M., Nakakura, T., Ebara, M., Tomura, H., Mogi, C., Im, D.S., Murata, N., Kuwabara, A., Ito, S., Fukuda, H., Arisawa, M., Shuto, S., Nakaya, M., Kurose, H., Sato, K., Okajima, F., 2015. Characterization of Imidazopyridine Compounds as Negative Allosteric Modulators of Proton-Sensing GPR4 in Extracellular Acidification-Induced Responses. PLoS One 10, e0129334.

Ulyanova, T., Scott, L.M., Priestley, G.V., Jiang, Y., Nakamoto, B., Koni, P.A., Papayannopoulou, T., 2005. VCAM-1 expression in adult hematopoietic and nonhematopoietic cells is controlled by tissue-inductive signals and reflects their developmental origin. Blood 106, 86–94.

Vainer, B., Nielsen, O.H., Horn, T., 1998. Expression of E-selectin, sialyl Lewis X, and macrophage inflammatory protein-1alpha by colonic epithelial cells in ulcerative colitis. Dig Dis Sci 43, 596–608.

Velcicky, J., Miltz, W., Oberhauser, B., Orain, D., Vaupel, A., Weigand, K., Dawson King, J., Littlewood-Evans, A., Nash, M., Feifel, R., Loetscher, P., 2017. Development of Selective, Orally Active GPR4 Antagonists with Modulatory Effects on Nociception, Inflammation, and Angiogenesis. J Med Chem 60, 3672–3683.

Wang, Y., de Valliere, C., Imenez Silva, P.H., Leonardi, I., Gruber, S., Gerstgrasser, A., Melhem, H., Weber, A., Leucht, K., Wolfram, L., Hausmann, M., Krieg, C., Thomasson, K., Boyman, O., Frey-Wagner, I., Rogler, G., Wagner, C.A., 2018. The Proton-activated Receptor GPR4 Modulates Intestinal Inflammation. J Crohns Colitis 12, 355–368.

Yang, L.V., Radu, C.G., Roy, M., Lee, S., McLaughlin, J., Teitell, M.A., Iruela-Arispe, M.L., Witte, O.N., 2007. Vascular abnormalities in mice deficient for the G protein-coupled receptor GPR4 that functions as a pH sensor. Mol Cell Biol 27, 1334–1347.

